# Sensory and choice responses in MT distinct from motion encoding

**DOI:** 10.1101/2021.06.24.449836

**Authors:** Aaron J Levi, Yuan Zhao, Il Memming Park, Alexander C Huk

## Abstract

Macaque area MT is well known for its visual motion selectivity and relevance to motion perception, but the possibility of it also reflecting non-sensory functions has largely been ignored. Manipulating subjects’ temporal evidence weighting revealed multiple components of MT responses that were, surprisingly, not interpretable as behaviorally-relevant modulations of motion encoding, nor as consequences of readout of motion direction. MT’s time-varying motion-driven responses were starkly changed by our strategic manipulation, but with timecourses opposite the subjects’ temporal weighting strategies. Furthermore, large choice-correlated signals were represented in population activity distinctly from motion responses (even after the stimulus) with multiple phases that both lagged psychophysical readout and preceded motor responses. These results reveal multiple cognitive contributions to MT responses that are task-related but not functionally relevant to encoding or decoding of motion for psychophysical direction discrimination, calling into question its nature as a simple sensory area.

## Introduction

Primate area MT plays a critical role in the perception of visual motion. A long line of study has established that MT’s encoding of motion direction is quantitatively consistent with perceptual sensitivity, that noise in its responses is correlated with behavioral variability, and that causal perturbations of its activity affect motion perception in lawful and substantial ways (***Newsome and Pare, 1988***; ***Britten et al., 1992***, ****1996****; ***Salzman et al., 1992***). Owing to this powerfully integrated set of findings, many models and experiments can safely assume that MT is the key place that the brain looks to for information about visual motion. However, these successes do not logically imply that MT only carries sensory information, leaving our understanding of MT at risk of overlooking additional signals and computations that are not aligned with representing motion for the sake of motion perception. In this work, we show that a manipulation of temporal strategy during motion discrimination reveals large signals in MT that are precisely related to components of performing the task, but which neither directly impact psychophysical performance nor reflect straightforward links between perceptual decisions and the sensory responses which informed them.

In addition to the large, classic body of work describing the form and fidelity of MT’s representation of visual motion (***Born and Bradley, 2005***; ***Cormack et al., 2017***), some prior work has identified cognitive modulations of MT’s sensory-driven activity. Such modulations are still interpretable with respect to MT’s representation of visual motion direction, however. Most notably, attention can modify the sensory-driven responses of MT neurons, typically boosting the gain of responses (***Treue and Maunsell, 1996***; ***Seidemann and Newsome, 1999***; ***Cook and Maunsell, 2004***). These modulations of stimulus-driven activity modify MT’s representation of motion, and thus play out in behavior as if the visual motion itself had been modified. In contrast, recent work has shown that MT’s choice-correlated activity is distinguishable at the population level from its sensory-driven responses, and follows a different time course than the read-out of motion, as inferred from the psychophysical behavior (***Zhao et al., 2020***). While this intriguing initial observation suggests the existence of task-related signals not directly related to motion encoding, interpretation of this choice-related activity is constrained by the lack of any direct experimental manipulation of the decision-making process.

To directly test for and characterize non-sensory signals in MT, we manipulated the time course of psychophysical weighting while monkeys performed a direction-discrimination task, coupled with ensemble recordings of multiple neurons in MT analyzed via population-coding techniques. We explicitly manipulated whether early or late parts of the stimulus had stronger or weaker motion evidence on average, which affected the time course of how the visual motion stimulus was weighted for task performance, as assessed via psychophysical reverse correlation. This manipulation of temporal weighting strategy provided critical interpretive leverage for distinguishing the time courses of decision formation and choice-correlated activity, and also caused a surprising and strong modulation of the sensory responses themselves that was also not directly related to forming decisions about motion.

When perceptual weighting was unconstrained, direction-discrimination behavior was based primarily on early portions of the stimulus, the sensory representation showed a standard and modest falloff over the course of stimulus presentation, and a distinct and substantial choice-correlated response emerged during late portions of stimulus viewing. When we shifted the temporal readout strategy to favor late portions of the stimulus, behavior relied preferentially on later portions of the stimulus, but *later* portions of the sensory response were *decreased*, as opposed to increased. Choice-correlated activity was significantly muted during the late-weighting condition. However, choice-correlated activity was present after the stimulus, leading up to the response (a novel phenomenon evident across all strategic conditions, in fact). When subjects’ temporal weighting strategy was then manipulated to preferentially rely on earlier portions of the stimulus, later portions of the sensory response were increased, and choice-correlated activity was again evident during the late portions of the stimulus. This last condition’s effects were most striking, as a steep falloff in perceptual weighting over time was accompanied by an increase in late sensory-driven activity that led to a non-monotonic time course of motion-driven response.

The opposite effects of our experimental manipulations on temporal weighting strategy and the timecourse of sensory gain run counter to any standard encoding model of MT simply representing behaviorally-relevant motion: In that framework, motion responses ought to mirror the psychophysical weighting. Choice-correlated activity during the stimulus was also controlled by changes in the psychophysical weighting, and across these psychophysical time courses, was always lagged relative to the periods when the subjects were “reading out” MT activity. But this decision-lagged choice-related signal was not simple feedback linking a sensory response and a subsequent, corresponding decision, not just because the choice signals affected MT population activity differently than visual motion did; we also observed a distinct choice-related signal after stimulus offset that was linked to impending response, and which was distinct from simple premotor activity.

Together, these multiple components of the MT response, revealed while manipulating the temporal weighting strategy, could be seen as lawful functions of the time course of decision formation and the anticipation of the response. However, these response components could not be interpreted as either modulations of the encoding that played out in perceptual reports, nor to the effects of read out mechanisms that would either correlationally (via feed-forward mechanisms) or causally (via straightforward feedback mechanisms) align with the sensory response. Thus, there appear to be multiple, large components of MT activity that affect both its stimulus-driven activity and which are separable from it– even during a well-studied direction-discrimination task with tight control over motion readout strategy– that are inconsistent with its conventional designation as a simple, low-dimensional, sensory encoding area.

## Results

We measured the timecourse of sensory and choice-correlated responses from simultaneously recorded groups of MT neurons using linear and nonlinear decoding approaches while rhesus monkeys performed a motion direction discrimination task. We manipulated the time course of stimulus evidence, and the subjects shifted their temporal weighting strategy to rely preferentially on the stronger periods of stimulus motion. We began recordings in each subject with a baseline “flat” stimulus phase for several experimental sessions, in which stimuli had a constant average motion strength over time within a trial, as is the case in most related experiments (***Gold and Shadlen, 2007***). We then shifted to several sessions in a “late” regime, in which the stronger motion was present in the second half of the stimulus. Finally, we performed several sessions in an “early” regime, in which the stronger motion was present in the first half.

### Observers change temporal weighting strategies according to stimulus statistics

Two trained rhesus macaques (one male, one female) viewed sequences of seven motion pulses and indicated perceived net motion with a saccade to one of two response targets (Figure 1A). We measured traditional psychometric performance (i.e., accuracy as a function of net motion strength on each trial), and the time course of weighting within each trial (i.e., using logistic regression between motion strength at each pulse and the binary choices, see Methods). We refer to the resulting set of regression coefficients, or weights, as the temporal weighting strategy.

**Figure 1.**
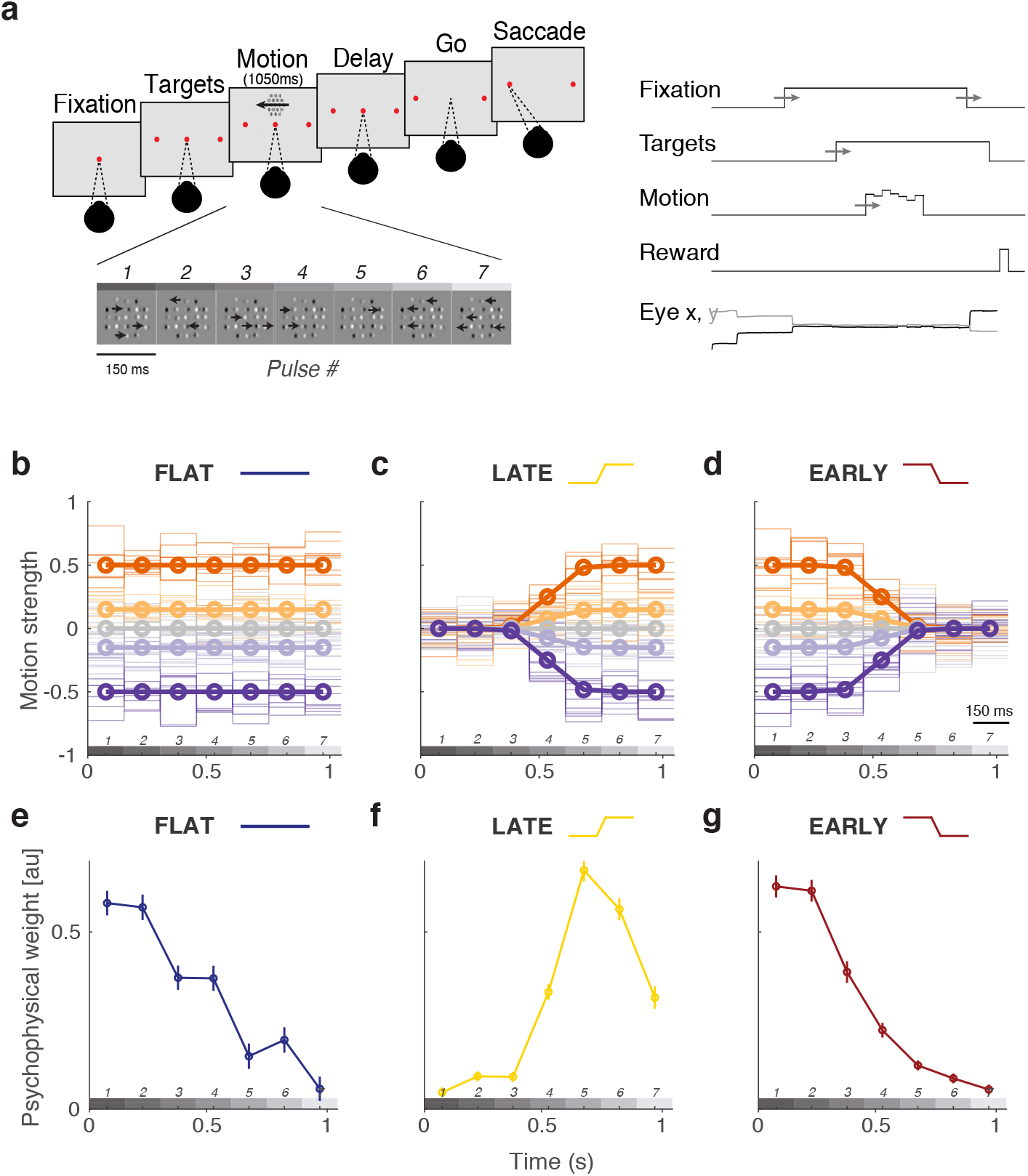
Sequence of trial events, temporal stimulus statistics, and successful manipulation of behavioral weighting strategy. **A**, Subjects fixated on a central point through the appearance of targets and motion stimulus until the disappearance of the fixation point (“go”). Choices were made with saccades to the target corresponding to the perceived net direction of motion. Initial fixation time, target-on duration, and time until fixation point disappearance were randomly varied. **B-D**, Average stimulus strength per pulse (bold lines) and individual trial examples (semi-transparent lines) for trials of different strength and direction (denoted by sign). In the flat-stimulus **(B)**, motion strength is constant over time on average. In the late-stimulus **(C)** motion strength is reduced on average in the first three pulses such that the highest motion expectation is late. In the early-stimulus **(D)** motion strength is reduced in the last three pulses such that the highest motion expectation is early. Motion pulse values in individual trials (semitransparent traces) vary considerably (see Methods for detail). **E-G**, Temporal weighting behavior across conditions. **E.** Subjects preferentially weight the early pulses despite uniform motion expectation over time. **F.** Temporal weighting shifts during the late-stimulus condition to preferentially weight late pulses. **G.** Behavior reverts back to early-weighting when the stimulus statistics are biased towards high motion strength early.

The motion discrimination task was performed in three contexts (Figure 1B-D). First, in the flat-stimulus condition (Figure 1B), average motion over time was equal within a trial. Many traditional sensory decision-making studies use stimuli with uniform information over time, and thus the flat-stimulus condition served as a baseline in our experiments. Subjects’ temporal weighting strategies were biased to have higher weight on early stimulus periods, despite uniform motion expectation over time (Figure 1E). This default early weighting strategy is consistent with many other findings (***Huk and Shadlen, 2005***; ***Kiani et al., 2008***; ***Nienborg and Cumming, 2009***; ***Yates et al., 2017***; ***Levi et al., 2018***; ***Kawaguchi et al., 2018***) and likely reflects a combination (***Levi and Huk, 2020***; ***Okazawa et al., 2018***) of improved sensory encoding at stimulus onset (***Osborne et al., 2004***; ***Churchland et al., 2010***), and the consequences of early termination of the decision process, due to mechanisms like bounded accumulation (***Kiani et al., 2008***).

Next, we performed a series of experimental sessions in which the stimulus statistics were manipulated such that the average motion strength was high for the last three pulses, while the first three were near zero. We refer to this as the late-stimulus condition (Figure 1C). Although the first 3 pulses had motion strength near zero on average (regardless of full-trial, net motion strength), on individual trials there was still variable nonzero motion possible for any pulse. Subjects were rewarded based on the actual net motion direction presented on that particular trial, as opposed to the average or expected value based on the condition from which the trial was generated. This produced robust behavioral changes that tracked motion expectation in the stimulus design, such that weight on the first three pulses decreased substantially, and the highest psychophysical weight was placed on the later pulses (Figure 1F).

Finally, we performed a series of sessions in which the stimulus statistics were changed such that the average motion strength was now high in the early half of the stimulus, and was near zero for the last half of the stimulus; we refer to this as the early-stimulus condition (Figure 1D). This successfully changed temporal weighting behavior back to pronounced early weighting, in which the first pulses received drastically higher weight than the remainder of the stimulus (Figure 1G), in a manner overall similar to the default strategy during the flat-stimulus (flat: −0.091 [−0.113, 0.069], late: 0.083 [0.015, 0.151], early: −0.091 [−0.136, −0.081]; slope of linear fit to the psychophysical kernel [95% CIs]). In summary, the temporal weighting strategy shifted in concert with the time course of expected motion strength, placing higher weight on portions of the stimulus when higher motion strength was expected based on the experimental phase. This confirms that our manipulation of stimulus statistics affected the time course of psychophysical readout, allowing us to better interpret the time scale of neural responses relative to the timing of when the subject was “reading out” MT for the purpose of forming a decision about motion direction.

### Choice-correlated activity in MT is large but does not align with stimulus encoding or behavioral readout

We recorded ensembles of single and multi-unit activity from area MT while monkeys performed the direction discrimination task, across the manipulation of temporal weighting strategy described in the previous section. We used both linear and nonlinear ensemble decoding frameworks to extract information about direction and choice from groups of simultaneously recorded MT neurons (Figure 2A). As a simple starting point, we used logistic regression (logReg) between the raw trial spike count vectors and either the stimulus direction (the “direction” axis) or the psychophysical choice (the “choice” axis) to find a reweighted population response that best mapped neural activity to the binary stimulus or choice (Figure 2A, left). Such linear models are likely easy for the brain to implement, but are limited in how they can capture relations between neurons and between neural activity and experimental factors. We therefore also used a more advanced nonlinear dimensionality reduction technique (variational latent Gaussian process model, vLGP) to extract smooth low-dimensional latent factors that explain correlations within the population spike trains (***Zhao and Park, 2017***; ***Zhao et al., 2020***) (Figure 2A, right). It functions in a conceptually analogous manner to the simpler logistic regression approach (i.e., mapping ensemble activity to the stimulus or the choice), but has the ability to more effectively capture the complex joint statistics of the neural population while also providing access to a more concise representation of the relations between neural activity, stimulus direction, and psychophysical choices (by virtue of an intervening dimensionality reduction step to identify latent factors making up the ensemble activity).

**Figure 2.**
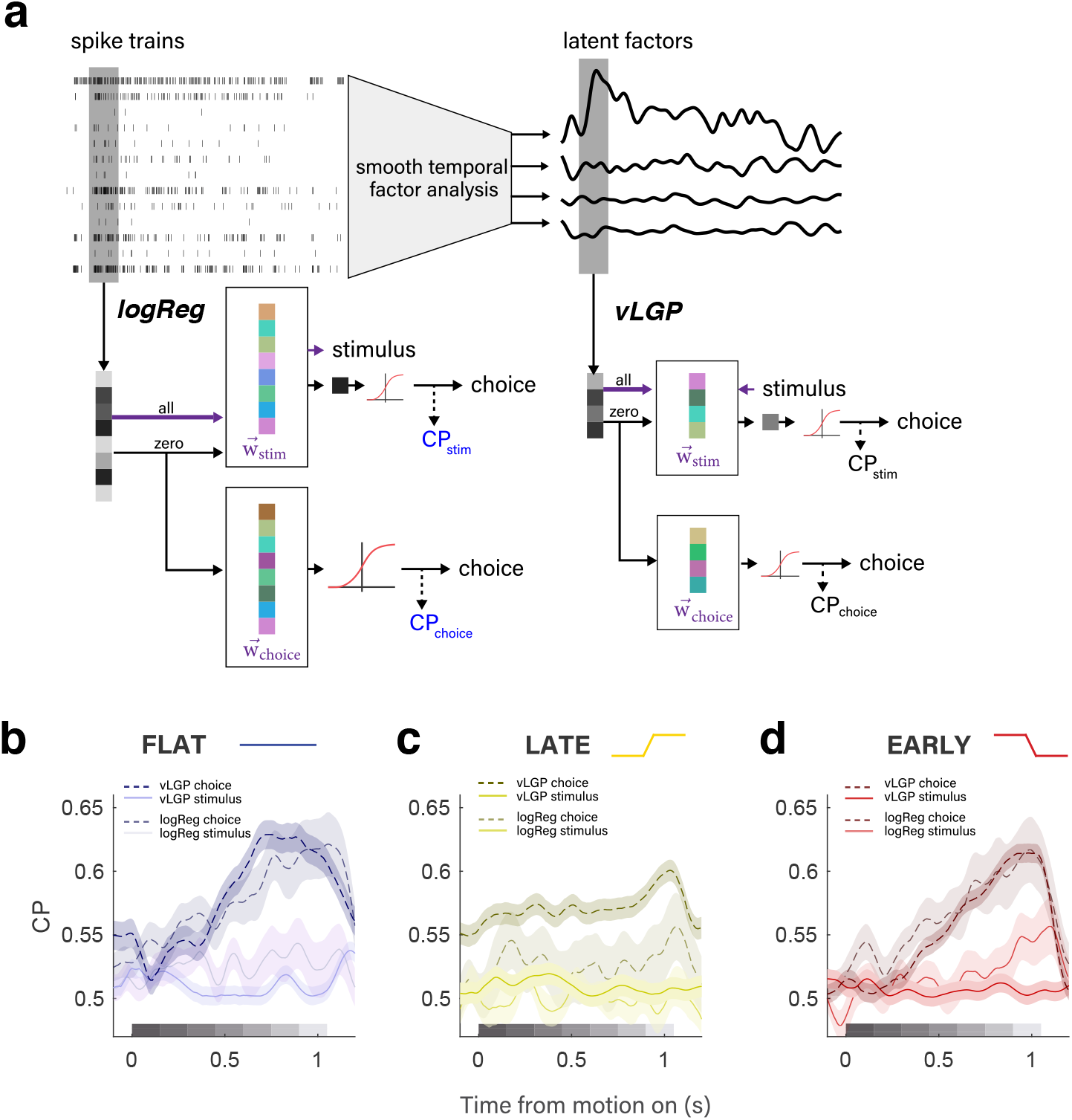
Both linear and nonlinear ensemble analysis approaches reveal strong choice-correlated activity in MT distinct from motion encoding or psychophysical readout of motion signals. **A.** We used linear and nonlinear decoding approaches to define choice probability along different dimensions of the population response. From the simultaneously recorded spike trains, a linear projection that can best predict the stimulus direction 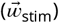 or the choice 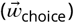 is used to project the frozen-noise trials and in turn derive CP (left). To enhance the signal to noise ratio, we extracted low-dimensional latent factors that explain the correlations in the population spike trains using smoothing factor analysis (right). We similarly estimated two CP signals from the latent factors. The first projection is found by the singular dimension explaining the stimulus drive for all trials 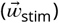. The second is the choice information extracted from the top four latent factors altogether 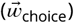. Projection of the frozen-noise trials are still multi-dimensional, and require further logistic regression to best predict the choice, defining the projection 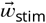 and corresponding choice probability CP_choice_. **B-D.** Time course of population choice probability during flat **B**, late **C**, and **D** early conditions. Solid vs. dashed line denote stimulus vs. choice dimensions, respectively. Darker traces in the foreground denote latent factors, while semi-transparent traces denote logistic regression traces in the background.

Both analytic approaches revealed the presence of substantial choice-correlated activity in the MT population response, achieving large peak magnitudes (> 0.6 as measured by choice probability, CP; although we use this conventional metric in this paper, we emphasize that by calculating it on various dimensions of the ensemble response, we have generalized it beyond the classical approach of only looking at choice-correlated activity defined along the stimulus axis) (***Britten et al., 1996***). The largest choice-correlated activity was present in the population activity in a manner distinct from how the stimulus drove the ensemble of MT neurons. Via logReg, this was evident in significantly larger CP along the choice axis over the direction axis (Figure 2B-D), stemming from a weak correspondence between a neuron’s weight in one model compared to the other (r = 0.146). The vLGP analysis showed that stimulus encoding was well described by a single dimension (termed the stimulus axis), but the stimulus axis had relatively small choice information when compared to the combined choice information in the top four latent factors altogether (***Zhao et al., 2020***) (Figure 2B-D).

Importantly, both analysis methods revealed that across pronounced changes in temporal weighting strategy, the time course of choice-correlated activities never mirrored the time course of psychophysical readout (Figure 2B-D, 1E-G). Instead, choice-correlated activity was consistently highest after the stimulus periods that were weighted the highest in the behavior. In the flat condition, both analysis approaches demonstrated increased choice probability during the last half of the stimulus, despite early weighting in the behavior. In the late condition, when behavior exhibited the strongest dependence on later portions of the stimulus, the strongest choice-correlated activity was still distinct from the stimulus-driven activity, and exhibited a more muted and flatter time course, though still characterized by an even later peak relative to the flat-stimulus condition. Finally, when subjects returned to an early weighting strategy in the early stimulus condition, the time course of choice probability returned to a similar rising profile, as originally measured during the flat condition. These observations are inconsistent both with classical interpretations that choice probabilities reflect the feedforward consequences of sensory noise being read out as information about the stimulus (because the bulk of the choice-correlated activity arose after the psychophysical readout of MT was likely happening), as well as more recent interpretations that choice probabilities reflect feedback, because differential MT responses correlated with choice were not strongly aligned with the motion responses that gave rise to those decisions.

### Changes in sensory encoding run opposite changes in temporal weighting strategy

Most surprisingly, we observed large changes to MT’s time-varying sensory response that were incommensurate with perceptual readout. Here, we relied on the vLGP analysis to describe the temporal dependence of the population response on the motion pulses by looking at the directional response along the stimulus axis. We calculated a pulse-triggered average (PTA) to summarize the regression coeffcients that predict the change in latent states (***Yates et al., 2017***). Each “bump” in Figure 3 represents the modulation of the stimulus-axis latent factor by a pulse of unit motion strength (i.e, a single Gabor drifting in one direction) for each pulse in the stimulus presentation (Figure 1). As temporal weighting strategy shifted across conditions, one might expect nothing to change in MT, consistent with a constant (and thus largely veridical) representation of visual information despite changes in readout/weighting strategy. An alternative hypothesis based on temporal attention would predict gain modulation congruent with behaviorally up-weighted and down-weighted stimulus epochs (***Ghose and Maunsell, 2002***). Instead, to our surprise, we observed changes to sensory encoding with an unintuitive, if almost paradoxical, link to psychophysical direction discrimination.

**Figure 3.**
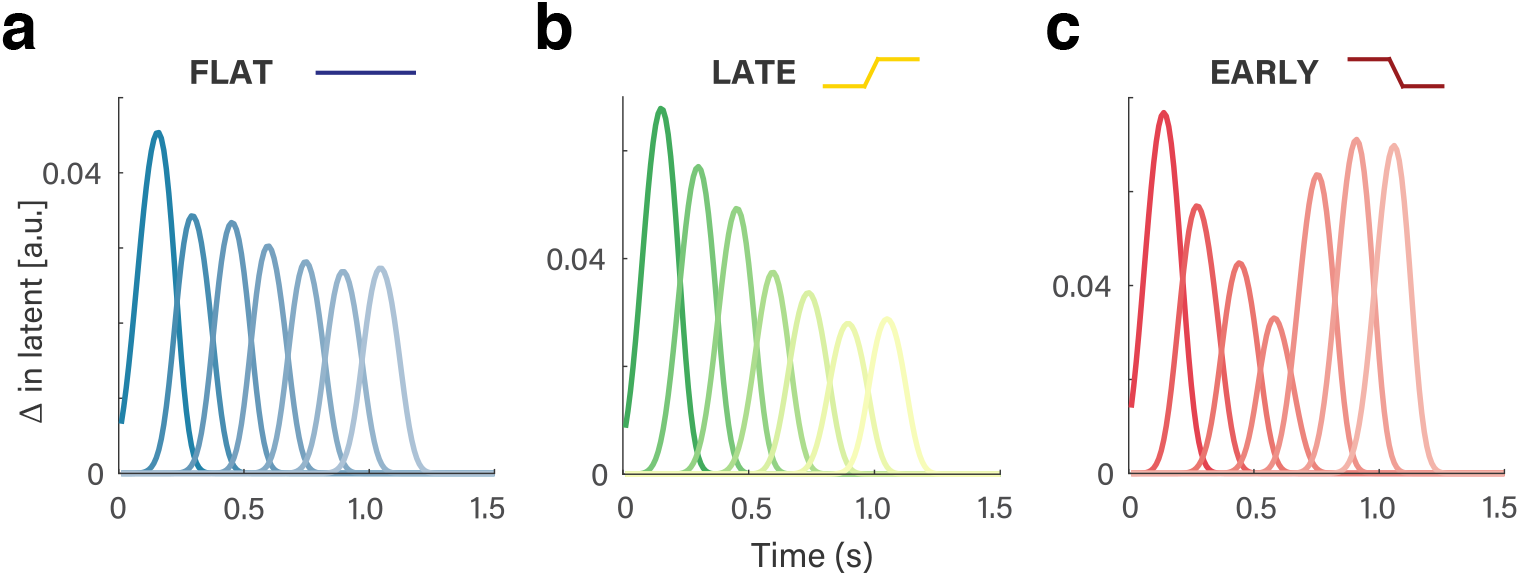
Time course of motion-driven MT response changes opposite that of changes in temporal weighting strategy. **A-C.** The pulse-triggered average (PTA) describes the modulation of the stimulus-axis latent factor by a pulse of unit motion strength for each of the seven pulses in the visual motion stimulus. **A.** The PTA for the flat-stimulus condition reflects the expected transient-to-sustained response, where a pulse at the beginning of the stimulus affects the MT response more than a pulse closer to the end of the stimulus. **B.** In the late-stimulus condition, the relative drop from early pulses to later ones is even more exaggerated than in the flat, despite highest motion strength occurring late in the trial. **C.** The PTA during the early-stimulus condition exhibits substantial increase on later pulses, despite a lack of high motion signal in the stimulus during those pulses.

In the flat stimulus condition there was a modest decrease in the sensory response over time (i.e., PTA magnitude fell across the 7 pulse epochs; Figure 3A). Such a gradually-declining time course is consistent with known adaptation phenomena in many visual brain areas, and has been observed in MT during viewing of this same stimulus (***Yates et al., 2017***). However, during the late-stimulus condition, the sensory response decreased for the late pulses relative to the flat condition time course (Figure 3B). The behavioral profile shows precisely the opposite: relative down-weighting of early pulses and up-weighting of later pulses. And most strikingly, when subjects switched to the early-stimulus condition, the sensory response showed a stark up-weighting of later pulses, resulting in a dramatically non-monotonic, U-shaped profile (Figure 3C). Once again, this is directly at odds with the temporal weighting of behavior, which sharply favors the first 2-3 pulses over the rest. This modulation is counterintuitive from standard perspectives, which would predict that if any changes in sensory response are evident, they would be reflected by increases in response to stimulus portions that were weighted more strongly for decision making.

Instead of gain changes that reflect behavioral readout strategy, the sensory response modulations we observed make more sense viewed as compensating for “missing” signal relative to a time-stationary motion expectation. In our experiments, both animals were trained extensively on the flat condition before undergoing temporal manipulation. The change in gain thus manifested as a function of the mismatch between this apparently “default” temporally-uniform expectation of motion and the statistics of the currently-encountered condition. In more detail, during the late condition motion strength was decreased in the early portions of the stimulus, but the PTA revealed decreased gain on later pulses instead (Figure 3B). During the early condition, the motion strength on later pulses was decreased, but the PTA revealed a striking gain increase on these portions of the stimulus for which the expected motion was quite weak (Figure 3C). Thus, while the temporal weighting evident in behavior changed across conditions in a way that tracked changes in stimulus statistics (i.e., weighting the stronger periods of motion more, and weaker periods of motion less), MT’s response to motion was changed inversely to those patterns.

### Large choice-correlated activity also exists in the absence of the motion stimulus

We also observed another choice-related signal in MT of substantial magnitude. The vLGP analysis revealed significant choice-correlated activity after the offset of the motion stimulus, in anticipation of an upcoming saccade. There was a minimum 500 ms window between the stimulus offset and the disappearance of the fixation point which signaled the monkey could move their eyes to make their choice, and during this window we saw choice probabilities up to > 0.7 (Figure 4A).

**Figure 4.**
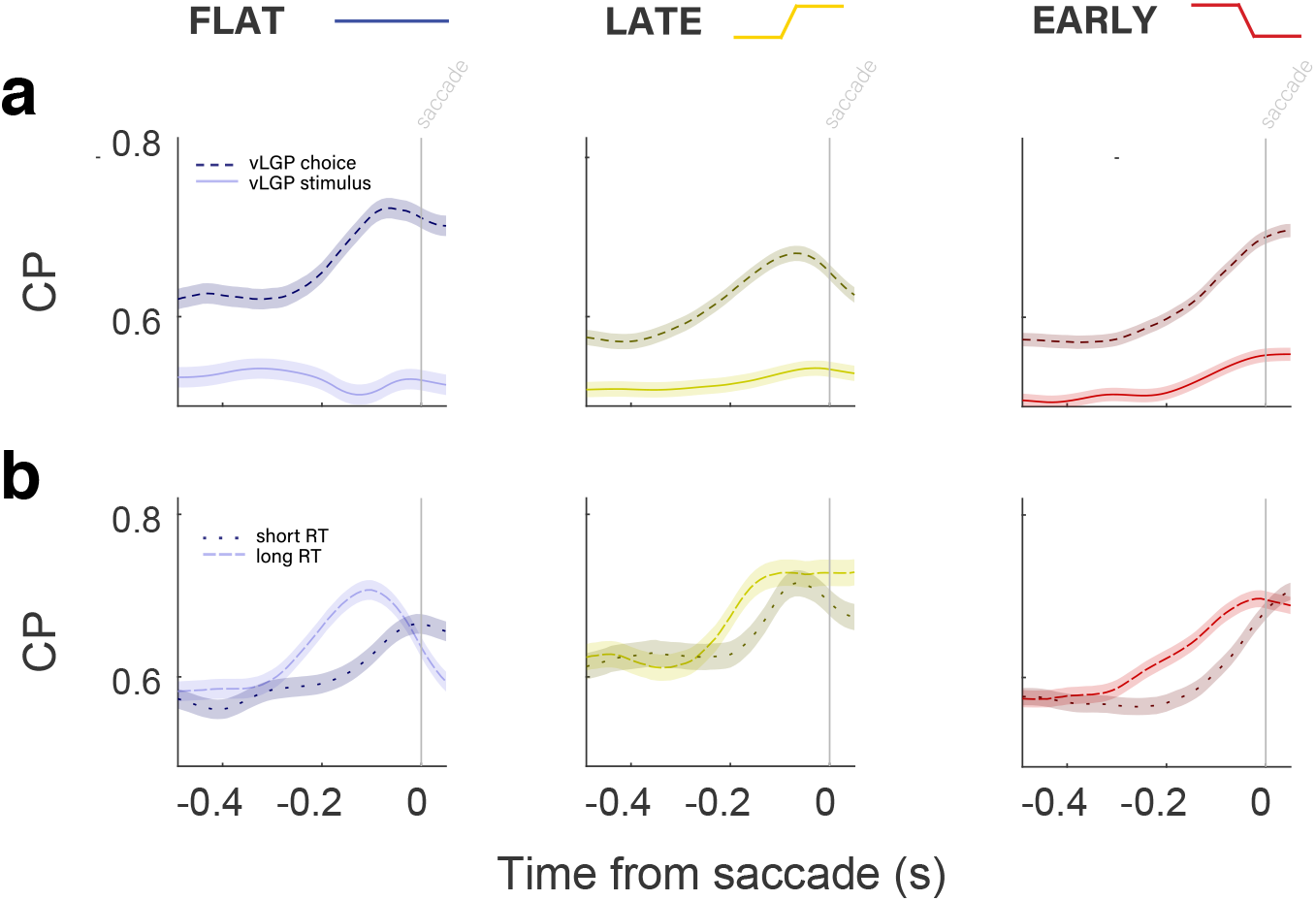
Presence of large choice-related signals in MT during post-stimulus delay period. **A.** CP along the choice (dashed lines) and direction (solid lines) axes, aligned to the time of the saccade. In all three conditions, there is high CP in the choice axis for the entire 500ms preceding the saccade, without any stimulus drive. CP increased over the last 200ms leading up to the saccade. There was realtively little CP along the stimulus axis. **B.** Saccade-aligned CP along the choice axis only, separated by median reaction time (RT). CP for longer RT trials (dashed lines) increased earlier than that of shorter RT trials (dotted lines). This was true in all three conditions.

The magnitude of post-stimulus choice probability is comparable to, and often greater than, what we observed from our decoders during the stimulus period, and is quite high compared to traditional measures of choice probability based on single neuron measurements. Most importantly, the finding of large amounts of choice-correlated activity without the presence of a visual stimulus in MT strengthens the case for such signals being non-sensory in origin. The choice signal measured during the delay period is present when there is no sensory drive whatsoever, further ruling out interpretations of choice probabilities as a product of noise in sensory representations. Instead, its full magnitude (revealed by “looking” off the stimulus axis), late time course, and presence up to the response are more similar to choice-related activity seen in a multitude of areas that are often considered much more cognitive or associative in nature, such as LIP and PFC (***Roitman and Shadlen, 2002***; ***Mante et al., 2013***).

Interestingly, the onset of CP during the delay period varies with reaction time (RT) in a way that suggests the choice signal is not simple premotor activity. If this were the case, we would expect that CP would increase later on trials with longer RTs compared to trials with shorter RTs. Instead, when reaction times were longer than the median RT, the saccade-aligned CP increased noticeably earlier than on trials with reaction times in the shorter half of the RT distribution (Figure 4B). This was true of all three temporal stimulus conditions. The result is striking, especially given the fixed-stimulus experimental design and the coarse division of “short” and “long” RTs by median. Temporally divorced from stimulus processing and not tightly linked to motor behavior timing, this delay-period choice signal appears to have a more cognitive origin reflecting the maintenance of choice information between stimulus and response.

### Time-varying readout of population activity confirms the dynamics of choice-related signals

In all analyses leading up to this point, the weights used to decode the stimulus or the choice were calculated using the neural responses and/or the derived latent factors from the entire stimulus period. Even with this fixed temporal readout scheme, we saw nuanced temporal dynamics in both sensory- and choice-related activity that differed from the time course of temporal weighting evident in the psychophysical behavior. Although, from a decoding perspective, using temporal fixed weights makes for a readout process that the brain might find easier to implement, we know very little about how sophisticated the brain’s decoding machinery might be (and indeed, our ability to manipulate the timecourse of motion weighting suggests that temporally-static decoding is not a hard limit). Furthermore, from a purely statistical perspective, we were also motivated to consider decoding with a temporally dynamic readout scheme to confirm that the rich dynamics we observed were neither constrained nor distorted by the assumption of constant read-out weights. We therefore performed further latent factor analyses in which weights were fitted and applied based on the activity within individual 100 ms bins for both the delay and motion periods (Figure 5).

**Figure 5.**
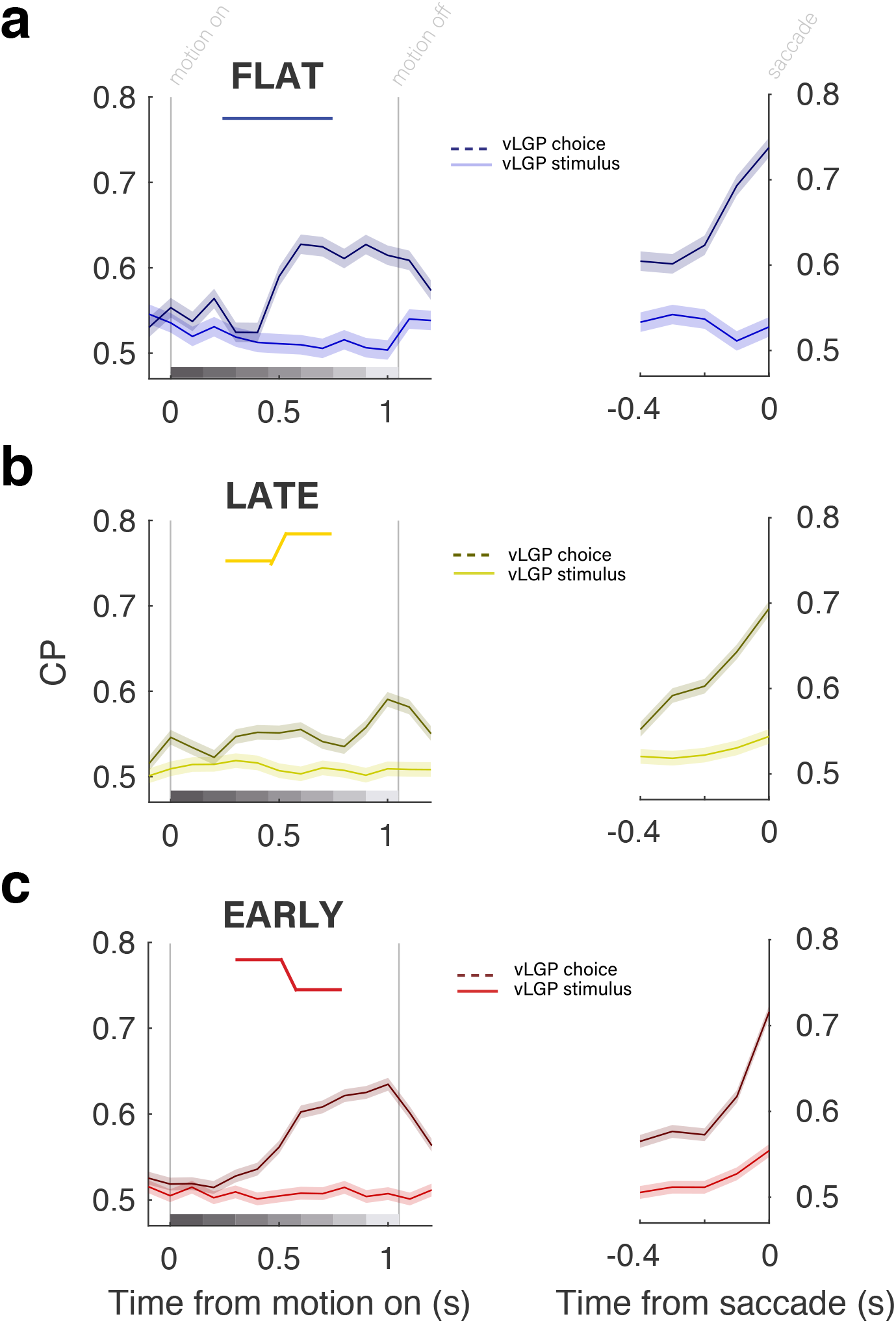
Time course of choice-related activity in MT is similar when time-varying decoding weights are used. Choice probabilities calculated with time-varying readout weights aligned to motion (left), and the saccade (right) for the flat (**A**), late (**B**), and early (**C**) conditions. CP along the choice axis is represented by dashed lines, while CP along the stimulus axis is represented by solid lines. Choice-axis CP was significantly higher in both the motion- and saccade-aligned time frames. During motion (left), we confirmed that CP was highest during later stimulus epochs, after those with highest psychophysical weight (Figure 1E-G.) During the post-stimulus period (right), we confirmed that CP increased primarily over the last 200ms preceding the saccade to levels even higher than motion-aligned CP.

The timecourse of choice-correlated activity was quite similar from fixed to dynamic readout models. With temporally varied readout weights, the same pattern persisted: high CP late in the stimulus period regardless of temporal stimulus condition. This is strong support for CP as a top-down signal that arrives in MT mostly after decisions have been made. That is, after the pulses with the highest weight in the psychophysical kernel. In this interpretation, during the late condition we have in essence delayed the decision and thus further delayed the decision-correlated activity that follows. The time-varying readout schemes also confirmed the dynamics in the post-stimulus, delay period. In all three conditions, CP was high throughout the delay period, but increased over the last 200ms. Along the stimulus axis, CP was flatter and closer to chance. Altogether, the similarity in CP timecourse between fixed and dynamic readout models suggests that a fixed weighting scheme is suffcient to describe the temporal patterns of choice information in MT during motion information both during and after the stimulus.

## Discussion

By manipulating the temporal weighting strategy of subjects while they performed a direction discrimination task, aided by ensemble recordings and population-level decoding analyses, we discovered multiple signals in MT that are distinct from its representation of motion direction, solidly established to be used by later decision stages for perceptual reports and behavior. Striking changes in sensory response were associated with the mismatch between the current strength of sensory evidence and prior, learned time courses of sensory evidence. Although these large modulations affected the sensory encoding, they appear not to have affected the psychophysical behavior. Choice-correlated activity was also surprisingly strong, but was delayed relative to temporal weighting behavior, even when the latter was under direct experimenter control. Furthermore, the choice-correlated activity was evident at the population level in a manner that was distinct from stimulus-driven responses in MT, and was “readout-irrelevant” as well, in that it was largest when the subjects were not primarily reading out the stimulus, or even viewing a stimulus at all.

The changes we observed in sensory responses may seem paradoxical at first, as the gain was increased for periods of the stimulus during which the subjects applied the smallest amount of weight in forming decisions. This is opposite the notion of attention affecting gain for parts of a stimulus that are more relevant for decisions (***Treue and Maunsell, 1996***; ***Seidemann and Newsome, 1999***). But, these modulations appear more sensible when viewed as resulting from a mismatch between trained statistics and the current ones. The hypo-responsivity to late pulses in the late condition, and the hyper-responsivity to those same late pulses during the early condition, could both reflect a compensatory response to motion in the current condition compared to the expectation of the temporally uniform stimulus on which animals were trained. Indeed, potentially-releated homeostatic mechanisms have been observed in sensory cortex (***Benucci et al., 2013***). Through this lens, the temporal changes in the PTA reflect a recalibration of incoming information to meet the expectation of a temporally-flat stimulus. Thus, even MT’s sensory responses are strongly affected by cognitive factors in ways that are dissociable from its well-established, but no longer sole role of representing retinal motion for the sake of perception and/or behavior.

Our findings regarding choice-related activity also add to the case for MT carrying substantial non-sensory signals. Having previously used ensemble recordings and population decoding to show that stimulus- and choice-related activity in MT are distinguishable (***Zhao et al., 2020***), our findings in this study add several important facets. First, we exerted explicit control over the time course of psychophysical weighting, which allowed us to experimentally dissociate the psychophysical weighting from the time course of choice-correlated activity. By shifting the temporal weighting strategy, we effectively changed the average time of the decision, allowing us to confirm that choice signals followed primary decision formation when under explicit experimenter control. Second, we saw choice activity of substantial magnitude during the post-stimulus delay period. This result rejects virtually any stimulus-based interpretation, as the choice signal was present when the sensory stimulus was not. These results also rule out straightforward forms of feedback creating choice-related activity, as those explanations require the decision-related feedback to be aligned with the sensory responses that gave rise to the corresponding choice. Furthermore, the delay period choice signal was not entirely explainable as premotor. Given all these distinctions, the oddly-parsimonious interpretation is that choice-related activity in MT is a distinct cognitive signal (or set of signals) that are best understood outside of MT’s encoding of visual motion. Although the presence of large choice-related signals in macaque MT may be surprising at first, recent work in other species (but also using ensemble recordings and analyses) has revealed widespread representations of choice and other task-related signals (***Musall et al., 2019***; ***Stringer et al., 2018***; ***Grün-demann et al., 2018***).

These findings provide new connections between MT function and well-established conceptual and empirical frameworks. The sensory modulations associated with mismatches between expected and observed timecourses of motion aligns with both predictive coding and reinforcement learning models, both of which are abstractly based on errors between expected and encountered elements within a task (***Rescorla and Wagner, 1972***; ***Engel et al., 2015***). Although our findings run opposite known effects of temporal attention (***Ghose and Maunsell, 2002***) or attention-related gating of sensory responses (***Seidemann et al., 1998***), some recent work has decoupled attentional modulations in MT and MST from task performance (***Recanzone and Wurtz, 2000***). Our dissociation between MT modulations and task performance may be related, although in our case, their dependence on the strategic history of the subjects revealed signals that are not wholly irrelevant to the task, but are just not related to the formation of decisions on a trial-by-trial basis. This opens up the possibility that some attention-like phenomena may arise from expectations of stimulus statistics, instead of being modulations of sensory data per se. The post-stimulus choice signals we observed in MT may be related to prior observations of small-amplitude, but tuned, persistent activity in MT (***Bisley et al., 2004***); our findings suggest that those initial observations of relatively small changes in spike rate may have simply caught a glimpse of larger non-sensory signals preceding the saccadic decisions mostly missed by single unit recordings that cannot see alternate effects on population activity across diversely-tuned neurons. Finally, related work using a motion categorization task has revealed strong non-sensory, category-related activity in area MST, but not area MT (***Freedman and Assad, 2006***; ***Zhou et al., 2020***). Such category-related activity can also be thought of as “choice-correlated”, as distinct from purely sensory-driven. Although the tasks, training histories, and analytic approaches differ between that work ours, our findings suggest that the apparent distinction between MT and MST regarding the presence of such category/choice activity might be less strict than previously observed. Again, the potential for ensemble recordings and corresponding ensemble analyses may have been critical for not just observing these non-sensory signals in MT, but for appreciating their substantial magnitude.

To conclude, our manipulation of temporal weighting strategy revealed a dissociation of sensory response gain from decision formation. Likewise, our approach of using ensemble recordings and population decoding allowed us to see large choice-related signals that were not just temporally dissociated from psychophysical weighting (or even stimulus viewing), but that were large in magnitude and distributed across the population in a manner distinct from how visual motion direction is represented. Together, these signals and modulations call for consideration of MT well beyond its role in encoding of retinal motion. Understanding the population coding structure and functional roles of such task-related but non-sensory computations are new open questions.

## Methods and Materials

### Stimulus presentation and design

Stimuli were presented using the Psychophysics Toolbox with Matlab (Math-works) using a Datapixx I/O box (Vpixx) for precise temporal registration (***Eastman and Huk, 2012***). Sample stimulus presentation code is available on request. Eye position was tracked using an Eyelink eye tracker (SR Research), sampled at 1 kHz. M1 was seated 57 cm away from a 150 cm × 86 cm rear-projection screen (IRUS; Draper Inc.) covering the central 106° × 73° of visual angle. Images were projected onto the screen by a PROPixx projector (VPixx Technologies Inc.) driven at a resolution of 1920 × 1080 pixels at 120 Hz. M2 viewed stimuli on a 55-inch LCD (LG) display (resolution = 1920 × 1080p, refresh rate = 60 Hz, background luminance = 26.49 cd/m2) that was corrected to have a linear gamma function. M2 viewed the stimulus from a distance of 118 cm (such that the screen width subtended 54° of visual angle, and each pixel subtended 0.0282° of visual angle). Auditory feedback was played at the end of every trial, and fluid reward was delivered through a computer-controlled solenoid.

Subjects were required to discriminate the net direction of a motion stimulus and communicate their decision with an eye movement to one of two targets, placed on either side of the motion stimulus. The sequence of task events is presented in Figure 1A. A trial began with the appearance of a fixation point. Once the subject acquired fixation and held for 750–1300 ms (uniform distribution), two targets appeared and remained visible until the end of the trial. 500–1000 ms after target onset, the motion stimulus was presented at a range of eccentricities from 4° to 12° for a duration of 1050 ms. The fixation point was extinguished 500–1000 ms after motion offset, and the subject was then required to shift their gaze toward one of the two targets within 600 ms (saccade end points within 3° of the target location were accepted). The timing of each event was randomly and independently jittered from trial to trial.

The reverse-correlation motion stimulus contained motion toward one direction or the opposite, with varying motion strength. Spatially, the stimulus consisted of a hexagonal grid of 19 Gabor elements, where individual Gabor elements were set to approximate the receptive field (RF) size of a V1 neuron, and the entire motion stimulus approximated the RF size of an MT neuron, which scaled based on eccentricity from fixation (***Van Essen et al., 1981***). Motion was presented by varying the phase of the sine-wave carrier of the Gabors. Each Gabor underwent a sinusoidal contrast modulation over time with independent random phase. Gabor spatial frequency (0.8 cycles/° sigma = 0.1 x eccentricity) and temporal frequency 5–6 Hz, yielding velocities of 5.55–6.66°/s, respectively) were selected to match the approximate sensitivity of MT neurons (***Bair and Movhshon, 2004***).

Each motion stimulus presentation consisted of seven consecutive motion pulses lasting 150 ms each (9 frames on the 60 Hz display, 18 on the 120 Hz display), producing a motion sequence of 1050 ms in duration in total. On any given pulse, a number of Gabor elements would have their carrier sine waves drift in unison to produce motion (“signal elements”), and the remaining would counter-phase flicker (“noise elements”). Within any given pulse, signal elements were spatially assigned at random within the grid, and all signal element drifted in the same direction.

Motion strength on pulse *i* was defined as the proportion of signal elements out of the total number of elements, the value of which was drawn from a Gaussian distribution, *X_i_ N*(*μ_k_, s*) and rounded to the nearest integer, where k is the distribution index for the five trial types (strong left, weak left, zero-mean, weak right, strong right). Thus, while each pulse within a sequence could take on any value (and either sign/direction) from distribution *N*(*μ_k_, s*), the expectation of a sequence would be *μ_k_* (Figure 1B-D). The subjects were rewarded for selecting the target consistent with the sign of the motion pulse sequence sum (i.e., the net direction), independent of the distribution *μ_k_* from which the pulses were drawn.

Subjects performed the motion-discrimination task with three variations of temporal stimulus statistics (***Levi et al., 2018***). First was the flat-stimulus, in which expected motion strength was uniform over time within a trial. In other words, the mean of the motion strength distribution *N*(*μ_k_, s*) would be held constant throughout a stimulus presentation. In other words, the mean of the distribution from which *X_i_* was drawn was fixed at (*μ_k_*), for pulses 1–7 (Figure 1B).

Next, subjects encountered the late-stimulus, where motion strength was reduced substantially in early pulses, but not late. In this condition, *μ_k_* is set to 0 for the first pulse (*i* = 1), and reaches its expected value (*μ_k_*) by pulse 7. Finally, the opposite is done for the “early-stimulus” condition (Figure 1D), in which the first pulses maintain mean motion strength equal to *μ_k_* and later pulses have a mean near zero. In the late- and early-stimulus conditions, the transition from *μ_k_* at pulse 1 to *μ_k_* at pulse 7 is governed by a logistic function with parameters chosen to result in a smooth transition between the first 3 and last 3 pulses (midpoint = 4, slope = 0.3).

All subjects began the experiments with the flat-stimulus condition (Monkey L: 13; Monkey N: 10 sessions). After multiple sessions of stable psychophysical performance, the stimulus was changed to the late-stimulus conditions (Monkey L: 11; Monkey N: 11 sessions). Finally, after multiple sessions of stable psychophysical performance the stimulus was changed to the early-stimulus condition (Monkey L: 11 sessions; Monkey N: 15 sessions). Subjects were exposed to only one stimulus condition per session and were not cued as to which stimulus condition they were viewing before or during any given session (other than the stimulus statistics themselves).

Throughout all conditions, there existed a subset of “zero-mean” trials in which *μ_k_* = 0 for all 7 pulses, regardless of whether the stimulus condition is flat, late, or early. Sessions also contained 5-10% frozen seed trials, which were identical stimulus displays. The “frozen noise” stimulus always summed to zero, had the same temporal structure across sessions, and was completely identical within sessions. Subjects were rewarded at random on frozen noise trials.

### Behavioral analysis

Subject choices in the direction-discrimination task were analyzed with a maximum likelihood fit of a three-parameter logistic function (***Wichmann and Hill, 2001***) assuming a Bernoulli distribution of binary choices, in which the probability of a rightward choice is p and leftward choice is 1 − *p*, where *p* is given by

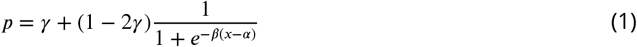

where *x* is the net motion strength value (z-scored over all sessions for each subject separately), a is the bias parameter (reflecting the midpoint of the function in units of motion strength), *b* is the slope (i.e., sensitivity, in units of log-odds per motion strength), and *g* captures the lapse rate as the offset from the 0 and 1 bounds. Error estimates on the parameters were obtained from the square root of the diagonal of the inverse Hessian (2nd derivative matrix) of the negative log-likelihood. The temporal weighting kernel (which we also refer to as “temporal weighting strategy” or “temporal weighting profile”) was computed using ridge regression via maximum likelihood. The log posterior of the psychophysical weights is given by

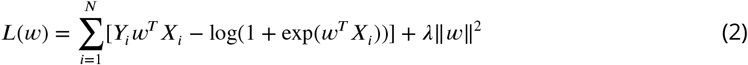

where *Y* ∈ {0, 1} is a vector of choice on every trial and X is a matrix of the seven pulses on each trial, augmented by a column of ones (to capture bias). *λ* was estimated using evidence optimization (***Sahani and Linden, 2003***). Psychophysical weights are normalized by the Euclidean norm of the vector of weights. The seven temporal weights assigned to the seven motion pulses, w, were computed by using all trials within a session. These include trials where *μ_k_* was set to zero (i.e. “zero-mean trials”, where motion on a given pulse is temporally independent of all other pulses in the sequence) and trials where *μ_k_* was set to a non-zero value (“signal trials”, where motion is correlated over pulses)

### Electrophysiology

A custom titanium chamber was fabricated and placed over the superior temporal sulcus and intraparietal sulcus to allow for a dorsal approach to access area MT. Chamber placement was as guided by structural MRI and cranial landmarks. Extracellular recordings were performed using linear electrode arrays from Plexon (U-Probe, V-Probe, or S-Probe; 24 or 32 channels; 50-100 micrometer spacing).

MT was identified using electrode depths and paths (i.e., sulcal anatomy), and functional mapping. Functionally, MT was identified via size and location of RFs, and preponderance of direction selective neurons. MT units were hand-mapped using a field of moving dots with experimenter control of stimulus location, aperture size, dot speed, dot size, and dot density. Upon choosing the stimulus location that maximally drove the highest number of neurons, direction tuning was measured by 500ms presentations of a randomly drawn direction of motion from one of 12 directions from 0 to 330 degrees. A total of 71 recording sessions were performed; 23 during the flat-stimulus condition (Monkey L: 13; Monkey N: 10), 22 during the late-stimulus condition (Monkey L: 11; Monkey N: 11) and 26 during the early-stimulus condition (Monkey L: 11; Monkey N: 15).

Spike sorting was performed using KiloSort (***Pachitariu et al., 2016***) followed by manual merging and splitting of clusters as necessary. A total of 583 units were identified; 161 during the flat-stimulus condition, 219 during the late-stimulus, and 203 during the early stimulus.

### Logistic regression neural decoder

To interrogate the roles and relationship of direction and decision-related signals, we used various decoding methods to approximate how information may be gleaned from groups of MT neurons. The first method we employed was logistic regression directly between spike counts and the binary direction or choice on each trial (***Kiani et al., 2014***; ***Yates et al., 2020***). The regression is done for each session such that each neuron is a feature in the model, where each neuron received a weight according to how well it predicts the binary outcome of interest. The result is a linear readout model that allows for maximal prediction of the stimulus direction or the animal’s choice.

Specifically, the decoding weights are calculated as coefficients in a logistic regression between trial spike counts (summed over a window starting at stimulus onset and ending 150ms after stimulus onset) and one of two binary variables (the stimulus direction, or the observer’s choice) using MATLAB’s glmfit. The choice decoder weights were calculated using only the zero-sum, frozen noise trials, while the direction decoder used all other trials.

The probability of a trial’s stimulus direction or choice given each neuron’s firing rate is given by:

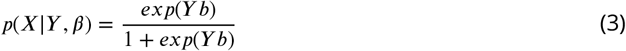

Where 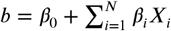 for *N* neurons present during a session. *X* is a vector of spike counts per neuron, and the choice or direction is *Y* ∈ 0, 1. The weights are then applied to their respective neuron’s temporally binned trial spike rates. Spikes were counted in 10ms bins and smoothed with a 50ms boxcar. This was expressed in terms of rates by dividing by the bin size. The result is a population response that best represented stimulus or choice information present in a recording session.

The resulting decoder output was then used to calculate population-level choice probability (CP) for each session. We measured CP over the course of stimulus presentation as a metric of trial-by-trial correlation between firing rate and choice, given a fixed stimulus. CP was calculated as the area under the ROC curve generated from choice-conditioned distributions of the reweighted activity in each temporal bin. CP time course traces were smoothed with a 100ms boxcar for visualization.

### Latent factor analysis

To understand how stimulus and perceptual choice are encoded across the population, we employed the variational latent Gaussian process (vLGP) method (***Zhao and Park, 2017***) to extract single-trial low-dimensional latent factors from population recordings in area MT. We used the recording between target onset and reward. The spike counts were binned at 10 ms. Let **x**_*k*_ denote the *k*-th dimension of the latent factors. We assumed that the spatial dimensions of latent factors are independent and imposed a Gaussian Process (GP) prior to the temporal correlation of each dimension,

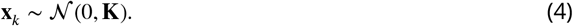

To obtain smoothness, we used the squared exponential covariance function and respective covariance matrix **K** in the case of discrete time. Let *y_tn_* denote the occurrence of a spike of the *n*th neuron at time *t, y_tn_* = 1 if there was a spike at time *t* and *y_tn_* = 0 otherwise at this time resolution. Then **y**_*t*_ is the vector of length *N*, total number of neurons in a session, that concatenates all neurons at time *t*. The spikes **y**_*t*_ are assumed to be a point-process generated by the latent state **x**_*t*_ at that time via a linear-nonlinear model,

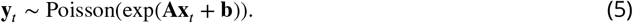

To infer the latent factors (**x**_*t*_ for each trial) and the model parameters (**A** and **b**), we used variational inference technique, as the pair of prior and likelihood do not have an tractable posterior. We assumed parametric variational posterior distribution of the latent factors,

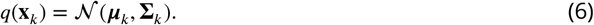

We analyzed the mean {***μ***_*k*_} as the latent factors in this study. The dimensionality of the latent factors was determined to be 4 by leave-one-neuron-out cross-validation on the session with the largest population. All the sessions with at least 4 simultaneously recorded units were included in this analysis (Monkey N: 13 sessions, Monkey L: 28 sessions).

#### Pulse-triggered average

To measure the relationship between the time-varying pulse strength and the inferred latent factors, we measured the contribution of pulses to the latent factors. The pulse-triggered average (PTA) measures the change in latent factors resulting from an additional pulse at a particular time of unit strength. To calculate the PTA, we used the pulse stimulus and latent response at 1 ms resolution. For each session, let *s_i_* denote the value of the *i*-th motion stimulus, and let *x_tk_* denote the *k*-th dimension of the latent factors at time *t*. All trials were concatenated such that the latent factors **X** is a matrix of length *T* × 4, where *T* is the total time. For the *i*-th pulse, *s_i_* is the number of Gabors pulsing, with *s_i_* > 0 for pulses in one direction and *s_i_* < 0 for pulses in the other direction. To calculate the temporal lags of the PTA, we built design matrices, **D** = [**D**_1_, **D**_2_, …, **D**_7_]. For the i-th pulse, the design matrix **D**_*i*_ is a *T* × 28 matrix that consists of 4 cosine basis functions at the 4*i* + 1, 4*i* + 2, …, 4*i* + 4-th columns and 0 elsewhere. These basis functions starts at 0 ms, 50 ms, 100 ms and 150 ms after the onset, lasts 100 ms each and spans the rows of **D**_*i*_. The magnitude of the bases is equal to the corresponding pulse value *s_i_*. We calculated a separate **D**_*i*_ for each of the seven pulses and concatenated them to obtain a design matrix for all seven pulses and estimated the weights with *ℓ*_2_-regularization,

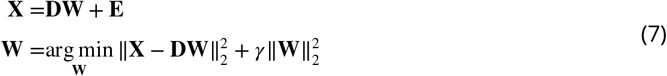

where **W** is the weight matrix to estimate and **E** is the Gaussian noise matrix and the regularization hyperparameter *γ* was chosen by the generalized cross-validation (GCV) (***Golub et al., 1979***). The PTA was calculated with the design matrices of unit-strength pulse and the estimated weights **W**. We smoothed the PTA with a temporal Gaussian kernel (40 ms kernel width).

Subject to arbitrary rotations, a latent trajectory forms an equivalence class of which the members have the same explanatory power in the vLGP model. We seek a particular rotation for each session that makes the encoded task signal concentrate in the first few dimensions. By singular value decomposition, **W**^⊤^ = **USV**^⊤^, we rotate the factors **x** to **U**^⊤^**x**.

#### Choice decoder

Since there were some recording sessions with less than ideal number of frozen trials (identical visual motion trials) for the calculation of choice probability, we instead analyzed the “weak” trials of which the monkeys’ correct rate was below a threshold (65%). We started at the trials of zero pulse coherence and gradually increased the magnitude of coherence (absolute value) until the correct rate reached the threshold. One of the sessions containing less than 100 weak trials was excluded in this analysis.

We removed the stimulus information that is encoded in the latent factors of weak trials by regressing out the pulses and analyzed the residuals. The latent factors were re-binned at 100 ms resolution where the value of each bin is the sum of latent state **x**_*t*_ or spike counts **y**_*t*_ over the bin for *t* = 1, 2, …, *T*. For each *t*, we assumed a linear model

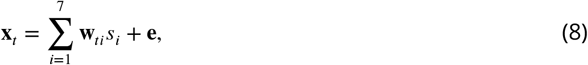

where *s_i_* denote the strength of the *i*-th pulse, **w**_*ti*_ is the weight vector corresponding to the bin and pulse, and **e** is the homogeneous Gaussian noise across all bins. We estimated the weight vector by least-squares with *ℓ*_2_-regularization to prevent over-fitting,

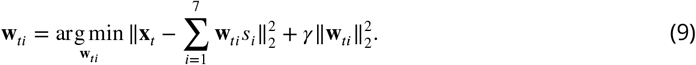

Again, the hyperparameter of regularization was chosen by GCV. We then analyzed the contribution of behavioral choice on the residuals

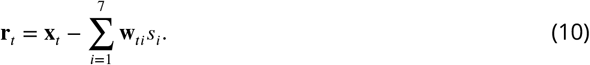

For the whole trial we used the sum residual of the windows **r** = **Σ**_*t*_**r**_*t*_. The range of t depends on the period of interest.

We trained logistic models, to which we refer to as *choice decoders*, to predict the choice on each trial using latent factors. The weights *β* and bias *β*_0_ were estimated by maximum likelihood with *ℓ*_2_-regularization,

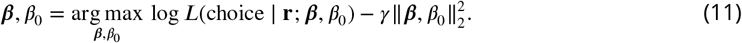

The hyperparameter of regularization was chosen via 5-fold (balanced classes in test set) cross-validation for every session individually.

#### Choice mapping

The conventional choice probability only applies to univariate variables. However, the latent factors and population activity are multivariate. We transformed the multivariate variables mentioned above onto a one-dimensional subspace that has the same direction as the choice through the choice decoders,

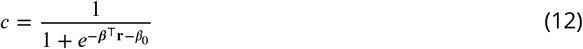

We refer to the transform as the *choice mapping*. The quantity *c* is a normalized value within [0, 1] that maps the residual onto the choice direction (***Lueckmann et al., 2018***), and enables pooling across sessions.

In order to prevent potential inflation of choice probability due to multidimensionality (3D), we regularized the choice decoder and used only the choice mapping on the test set (pooled samples held-out by cross-validation). This approach guarantees that choice probability will not be overestimated.

We pooled these mappings across all sessions. Using different subsets of latent factors as **r** in the mapping, we obtained the choice-mapping of the stimulus-dimension and non-stimulus-dimensions of latent factors. Then we calculated the choice probability of the corresponding dimensions based on the values. To investigate the time course of choice probabilities, we used choice decoders to perform choice-mapping on the whole dataset with non-overlapping moving windows. For fixed readout, we estimated the weights using mean value of 0-1.2s for the stimulus period and −0.5-0s for the delay period. We use the weights to obtain readout and CP values with 10ms moving window, and smooth the CP values with a 100ms boxcar. Finally; for dynamic readout, we estimated the weights and calculated the CP values within 100ms moving windows individually.

## Acknowledgments

Thanks to A Laudano, K Mitchell, and C Carter for animal support and C Badillo for technical support. I.M.P and A.C.H were supported by NSF IIS-1734910. A.J.L was supported by NIH/NEI R01-EY017366.

